# An Investigation into Mechanomyography for Signal Extraction and Classification of Human Lower Limb Activity

**DOI:** 10.1101/2024.12.01.626260

**Authors:** Yu Bai, Xiao Rong Guan, Rui Zhang, Shi Cheng, zheng Wang

**Affiliations:** the School of Mechanical Engineering, Nanjing University of Science and Technology, Nanjing 210000, China; Zhiyuan Research Institute, Hangzhou 310013, China

**Keywords:** Mechanomyography, Feature Mode Decomposition, Transformer model, Human Activity Classification

## Abstract

To mitigate the difficulties associated with the extraction of Mechanomyography (MMG) signals from raw Accelerometer (ACC) data and the subsequent classification of human lower limb activities based on MMG signals, the Feature Mode Decomposition (FMD) algorithm has been utilized for the isolation of the MMG signal. Simultaneously, surface Electromyography (sEMG) signals were recorded to perform correlation analyses, thereby validating the effectiveness of the extracted Mechanomyography (MMG) signals. The results demonstrate that the envelope entropy derived from the FMD was the lowest among the observed values, and the composite signal obtained via FMD displayed the highest correlation with the sEMG signal. This indicates that FMD is capable of efficiently isolating the MMG signal while maintaining the maximal quantity of muscle contraction data. To address the challenge of classifying human lower limb activities, a comprehensive feature extraction procedure was implemented, resulting in the derivation of 448 unique features from multi-channel mechanomyography (MMG) signals. Subsequently, Kernel Principal Component Analysis (KPCA) was employed to diminish the feature set’s dimensionality. This was succeeded by the deployment of a Temporal Convolutional Network integrated with an Attention mechanism (TCN-Attention) to train the classification model. Additionally, an enhanced Northern Goshawk Optimization Algorithm was leveraged for optimization purposes. The findings indicate that FMD exhibited the minimum envelope entropy value of 8.13, concurrently attaining the maximum correlation coefficient of 0.87 between MMG and sEMG signals. Significantly, the SCNGO-TCN-Attention model demonstrated superior classification accuracy, attaining an exceptional accuracy rate of 98.44%.

## 1. Note

A novel methodology for the extraction and categorization of human lower limb activities utilizing Mechanomyography (MMG) has been devised. This approach utilizes data acquired from Inertial Measurement Unit (IMU) sensors to facilitate the extraction of Mechanomyography (MMG) signals. The Feature Mode Decomposition (FMD) technique was utilized to isolate the Mechanomyogram (MMG) signal, whereas Kernel Principal Component Analysis (KPCA) was implemented for the preprocessing of MMG feature data. To classify human lower limb activities, the Temporal Convolutional Network integrated with an Attention Mechanism (TCN-Attention) algorithm was employed. Additionally, an enhanced version of the Northern Goshawk Optimization Algorithm was utilized for optimization purposes. The proposed extraction technique efficiently segregates MMG signals from acceleration datasets, while the classification model exhibits substantial promise for applications in human activity recognition. The present investigation encompassed human subjects, with all ethical considerations and experimental methodologies rigorously reviewed and sanctioned by the Medical Ethics Committee of Nanjing Medical University, as evidenced by approval identifier 2021-SR109.

## 2. Introduction

### 2.1. Introduction of Human Activity Recognition

Human Activity Recognition (HAR) has garnered significant scholarly attention due to its extensive utility across various domains, including healthcare, behavioral analytics, authentication systems, and fitness surveillance. Fundamentally, Human Activity Recognition (HAR) involves the systematic categorization of data streams originating from sensors into discrete movement and activity types, including but not limited to actions such as sitting, walking, and the utilization of tools. Typically, this objective is realized by employing inertial sensors integrated within devices such as smartphones and smart watches. The technology exhibits significant potential within medical settings, enabling enhanced patient monitoring, ensuring compliance with therapeutic protocols, mitigating unauthorized actions, and bolstering rehabilitation initiatives. Beyond the confines of the clinical domain, Human Activity Recognition (HAR) stands on the precipice of transforming interactions across diverse sectors, including gaming, robotics, and athletic endeavors. Despite notable advancements, numerous challenges remain that impede the extensive implementation of Human Activity Recognition (HAR) technologies, particularly concerning the assurance of their reliability within varied and uncontrolled environments. A significant challenge lies in the intrinsic variability observed in the execution of identical activities by distinct individuals, which is profoundly influenced by their unique personal attributes. The distinctiveness of individual patterns, although foundational for user identification, concurrently complicates the generalizability of activity classification models. The disparity is notably accentuated when models, initially trained on a specific cohort of subjects, are subsequently assessed on a distinct set, frequently resulting in a decrement in performance efficiency[1]. Amplifying the intricacies of this endeavor are supplementary elements including sensor-derived noise, the intricate computational demands of algorithms, and the inherently dynamic characteristics of motion patterns. These factors necessitate comprehensive mitigation strategies to augment the robustness and adaptive capabilities of Human Activity Recognition (HAR) systems.

### 2.2. Introduction of Transformer model in Human Activity Recognition

In the realm of human activity recognition (HAR), transformer models have emerged as potent instruments, yielding substantial enhancements in both accuracy and efficiency. Numerous pioneering frameworks have been introduced, each targeting distinct facets of Human Activity Recognition (HAR) through proprietary methodological approaches.

1. The ViT-ReT Framework: This integrated framework leverages the synergistic combination of visual and recurrent transformer architectures to achieve precise and computationally efficient Human Activity Recognition (HAR).The architecture is engineered to sustain superior accuracy while concurrently delivering substantial enhancements in processing speed compared to traditional Convolutional Neural Network (CNN) and Recurrent Neural Network (RNN) methodologies, thereby rendering it particularly suitable for deployment in resource-constrained and real-time application scenarios [2].
2. The Deep Reverse Transformer-Based Attention Mechanism employs a top-down feature reconstruction approach augmented with self-calibrated reverse attention. This configuration facilitates the efficient capture of both spatial and temporal characteristics, thereby enhancing the performance of sensor-driven Human Activity Recognition (HAR) systems. The method demonstrates superior performance compared to numerous contemporary state-of-the-art techniques across a range of publicly available datasets [3].
3. Convolution-Free Framework: This approach leverages Vision Transformers for frame-level feature extraction and Long Short-Term Memory (LSTM) networks for modeling long-range temporal dependencies. Consequently, it demonstrates superior accuracy on the UCF50 and HMDB51 benchmarks, thereby establishing novel benchmarks for video sequence-based Human Activity Recognition (HAR) [4].
4. The Convolution Visual Transformer Network (CVTN) embodies a synergistic integration of Convolutional Neural Networks (CNNs) and Visual Transformers, leveraging the former’s proficiency in spatial feature extraction and the latter’s adeptness in temporal pattern analysis. This integration enhances the precision and reliability of Human Activity Recognition (HAR) derived from sensor data, as evidenced by its superior performance on the Kinetics dataset [5].
5. The Human Activity Recognition Transformer (HART) is a specialized, lightweight transformer architecture tailored for the processing of inertial measurement unit (IMU) data collected from mobile devices. The model demonstrates exceptional performance in Human Activity Recognition (HAR) tasks, characterized by diminished computational demands and enhanced generalization capabilities across a variety of sensing contexts, thereby surpassing the efficacy of conventional Convolutional Neural Network (CNN) and Long Short-Term Memory (LSTM) architectures [6].
6. The Adapted Transformer Model for Motion Sensor Data employs the self-attention mechanism to effectively capture intra-time-series dependencies within smartphone-generated motion sensor data. This model demonstrates superior Human Activity Recognition (HAR) performance, achieving an accuracy of 99.2% on a substantial public dataset, thereby markedly surpassing the efficacy of traditional machine learning approaches [7].
7. The Transformer-Based Activity Recognition Model for Inertial Sensor Data presents a sophisticated and adaptable framework for extracting insights from inertial sensor datasets. This model consistently achieves superior accuracy and enhanced generalization across diverse datasets, encompassing extensive recordings from a multitude of users performing a wide range of activities. Its performance notably exceeds that of conventional Convolutional Neural Network (CNN) and Long Short-Term Memory (LSTM) architectures [8].
8. The Two-Stream Convolution Augmented Human Activity Transformer (THAT) Model: This model introduces an innovative dual-stream architecture coupled with multi-scale convolution-augmented transformers, designed to effectively capture both time-over-channel and channel-over-time characteristics from WiFi channel state information. This approach results in enhanced accuracy and efficiency in WiFi-based Human Activity Recognition (HAR) across various real-world datasets [9].
9. The Self-Attention Based Two-Stream Transformer Network (TTN) addresses the complexities associated with modeling spatial-temporal dependencies and processing multimodal sensor data. This is achieved through the deployment of distinct temporal and spatial streams, which are designed to extract complementary features, thereby enhancing the network’s ability to capture nuanced information from diverse data modalities. The system allocates attention weights to individual sensor axes in accordance with their respective contributions to the classification process, thereby exhibiting exceptional performance across a range of benchmark multimodal Human Activity Recognition (HAR) datasets [10].

The profound influence of transformer architectures on Human Activity Recognition (HAR) is unequivocal. These models leverage the self-attention mechanism to enhance both feature extraction and temporal modeling processes, resulting in substantial advancements in performance metrics. This advancement represents a pivotal milestone in the domain, establishing a foundational framework for the development of more advanced and dependable Human Activity Recognition (HAR) systems.

### 2.3. Introduction of Mechanomyography in Human Activity Recognition

In the domain of human activity recognition (HAR), mechanomyography (MMG) has been extensively utilized owing to its inherent versatility and demonstrated efficacy. The subsequent research underscores the potential efficacy of integrating Microbial Metagenomics (MMG) with sophisticated machine learning algorithms and multimodal sensor fusion techniques.

1. Facial Activity Recognition: Methodology: A comprehensive wearable sensor fusion apparatus, incorporating inertial, pressure, and acoustic mechanomyography (MMG) sensors, was seamlessly integrated into a sports cap to facilitate precise and dependable facial activity recognition, obviating the necessity for camera-based systems. Results: The system exhibited cross-user F1 scores reaching 82% in experiments involving 13 participants from varied cultural contexts, thereby showcasing an inclusive and privacy-preserving methodology [11].
2. Lower Limb Gait Movement Recognition: Methodology: A wireless mechanomyography (MMG) and attitude angle sensing system was engineered to discern 11 distinct lower limb gait movements, encompassing static postures, dynamic transitional phases, and dynamic activities. Utilizing Hidden Markov Models (HMMs), various feature selections, channel combinations, and assessments of muscle contributions were systematically evaluated. Results: The implemented system demonstrated an average recognition accuracy of 98.91% based on data derived from four thigh muscle groups [12].
3. Hand Movement Recognition: Methodology: An incremental learning approach leveraging deep neural networks was developed for the classification of four distinct hand movements, utilizing mechanomyography (MMG) signals acquired from eight distinct channels. Results: The proposed methodology demonstrated recognition accuracy rates reaching 88.25% when applied to non-identically distributed test datasets, surpassing the performance of multiple established machine learning algorithms [13].
4. Knee Motion Recognition: Methodology: A hybrid model integrating the automated feature extraction proficiency of Convolutional Neural Networks (CNNs) with the robust generalization attributes of Support Vector Machines (SVMs) was formulated to facilitate the recognition of knee motion through the analysis of 4-channel Mechanomyography (MMG) time series data. Results: The model demonstrated enhanced classification accuracy independent of manually engineered features, thereby rendering it particularly effective for application to small datasets [14].
5. Gesture Recognition: Methodology: A feed-forward neural network classifier was employed, utilizing features extracted from MMG signals acquired via five IMU sensors configured in a band arrangement. This approach integrated data pertaining to muscle activity and the relative orientation of the sensors. Results: The system exhibited an average F1 score of 94±6% across evaluations conducted on three subjects and three discrete orientations, thereby substantiating its efficacy in gesture recognition [15].
6. Knee Motion Intention Detection: Methodology: We developed a wearable mechanomyography (MMG) sensing system designed to ascertain knee motion intentions. This system employs a support vector machine (SVM) classifier, integrated with a Markov chain model and a novel feature extraction technique based on the difference of mean absolute values. Results: The system demonstrated a classification accuracy of up to 91% among a cohort of 8 participants, thereby establishing a foundation for the development of adaptable and ergonomically viable wearable human motion interfaces [16].
7. Hand Movement and Contraction Pattern Recognition: Methodology: Utilizing multi-channel mechanomyography (MMG) signals, the study aimed to discern hand movements and contraction patterns among both able-bodied individuals and amputee participants. The study utilized a genetic algorithm for feature selection, followed by linear discriminant analysis (LDA) for classification purposes. Results: The system demonstrated an accuracy rate of up to 90% in the identification of seven distinct hand movements, thereby enabling the precise estimation of relative muscle contributions and enhancing the comprehension of mechanical patterns associated with muscle activity [17].

The collective findings of these studies underscore the substantial promise of MMG-based sensing technologies in achieving high-precision recognition of human activities, movements, and gestures across diverse anatomical regions. Advanced machine learning paradigms coupled with multimodal sensor fusion methodologies significantly augment the performance and versatility of these systems, rendering them viable instruments across diverse application domains, including healthcare and rehabilitation, human-computer interaction, and other emergent fields..

### 2.4. Introduction of Research in Mechanomyography Extraction

In the domain of mechanomyography (MMG) signal extraction, a plethora of sophisticated signal processing algorithms have been devised to proficiently isolate pristine MMG signals from unprocessed data, thereby mitigating challenges associated with noise and motion artifacts.These methodologies employ advanced techniques such as Empirical Mode Decomposition (EMD), Variational Mode Decomposition (VMD), and their respective multivariate extensions, frequently integrated with optimization algorithms to enhance analytical precision.Several significant methodologies are worthy of mention:

1. Multivariate Empirical Mode Decomposition Augmented by Noise (NA-MEMD): Methodology: The Nonlinear Adaptive Multivariate Empirical Mode Decomposition (NA-MEMD) algorithm is specifically engineered to mitigate motion artifacts present in multichannel Mechanomyography (MMG) signals acquired during both isometric and dynamic muscle contraction phases.The methodology entails the decomposition of the signal into intrinsic mode functions (IMFs) to facilitate the identification and subsequent elimination of noise and artifact constituents. Results: The proposed methodology demonstrates superior performance compared to traditional filtering techniques, effectively mitigating artifacts and establishing a novel foundation for MMG feature extraction, frequency analysis, and crosstalk examination [18].
2. Multivariate Variational Mode Decomposition (MVMD): Methodology: The Multivariate Variational Mode Decomposition (MVMD) technique is employed to deconstruct multichannel Mechanomyogram (MMG) signals into bandwidth-constrained intrinsic mode function (IMF) components.The system effectively mitigates motion artifacts and noise through a selective summation process, which evaluates and integrates components based on their frequency and energy distribution characteristics. Results: The Multivariate Variational Mode Decomposition (MVMD) enhances signal recognizability and optimizes the energy of the Mechanomyography (MMG) signal, thereby surpassing the performance of traditional Multivariate Empirical Mode Decomposition (MEMD) methodologies [19].
3. Integrated Method Combining Multiple Techniques: Methodology: This method integrates several techniques: Application of Recursive Least Squares (RLS) Filtering for the Mitigation of Power Line Interference The Enhanced Gray Wolf Optimizer (EGWO)-Based Variable Mode Decomposition (VMD) Approach for the Segmentation of MMG Signals into Distinct Low and High-Frequency Bands The Complete Ensemble Empirical Mode Decomposition with Adaptive Noise (CEEMDAN) technique is employed for the precise extraction of intrinsic mode functions, utilizing predefined thresholds based on center frequency and sample entropy to ensure the isolation of significant components. Signal Reconstruction: The objective is to accurately reconstruct the mechanomyogram (MMG) signal subsequent to the implementation of noise suppression and artifact elimination protocols. Results: The integrated methodology demonstrates a significant reduction in noise and artifacts, concurrently maintaining essential MMG signal constituents, thereby circumventing the constraints inherent in the standalone VMD and CEEMDAN techniques [20].

These investigations underscore the efficacy of sophisticated signal decomposition methodologies and optimization algorithms in the precise denoising of raw mechanomyogram (MMG) signals, thereby enhancing the dependability and precision of MMG-driven human activity recognition and analytical processes.

### 2.5. Introduction of This Research

The majority of studies investigating the recognition of human movement intentions through the application of the Transformer model predominantly leverage image recognition techniques to accomplish motion classification tasks. Despite the initial promise demonstrated by these methodologies, they are accompanied by substantial limitations.

1. Controlled Settings: Image recognition methodologies typically necessitate controlled settings to guarantee the uniformity and superior quality of data acquisition.
2. Specialized Apparatus: The utilization of high-resolution imaging devices and tailored illumination configurations is generally requisite, presenting significant financial implications and often posing practical challenges in real-world deployment scenarios.
3. Computational Infrastructure: The processing of extensive image datasets necessitates significant computational capacity, presenting a potential impediment to broad implementation.
4. Controlled Environments: The necessity for regulated settings restricts the generalizability of these methodologies to real-world, uncontrolled contexts.
5. Privacy Considerations: The deployment of cameras for the purpose of capturing human movements engenders significant privacy issues, particularly within contexts that are deemed sensitive or inherently personal.

To mitigate these constraints, the present investigation centers on the utilization of mechanomyography (MMG) signals that antecede human motor activity for the purpose of discerning movement intentions. The principal elements of this methodology encompass:

1. Usability: The acquisition of MMG signals is facilitated through the utilization of wearable sensors, characterized by their portability, non-invasiveness, and adaptability for deployment across diverse environments.
2. Economically Prudent: Wearable sensors typically incur lower costs and obviate the necessity for specialized equipment that is requisite for image recognition technologies.
3. Real-World Applications: The deployment of wearable sensors enables the acquisition of data within natural, uncontrolled settings, thereby broadening the spectrum of applicable scenarios.
4. In contrast to prior investigations, the present study encompasses an extensive assessment of the quality associated with the acquisition of muscle tone signals. This entails evaluating parameters including envelope entropy and the correlation coefficient between mechanomyography (MMG) and surface electromyography (sEMG).

This study investigates the extraction and analysis of mechanomyography (MMG) signals through the application of the Feature Mode Decomposition (FMD) algorithm. Additionally, it explores the classification of lower limb movements utilizing a hybrid model that integrates the Sine-Cosine Northern Goshawk Optimization (SCNGO) algorithm with a Temporal Convolutional Network incorporating an Attention mechanism (TCN-Attention). The details are presented as follows.

MMG Signal Extraction and Analysis:

1. Algorithm for MMG Signal Extraction: Methodology: The Muscle Mechanomyography (MMG) signals elicited by muscle contractions were decomposed utilizing the Feature Mode Decomposition (FMD) algorithm [21].In addition to its utility in MMG signal processing, FMD has demonstrated efficacy across multiple domains, such as multi-sensor fault diagnosis, bearing fault diagnosis, and the detection of multiple localized faults within rotating machinery [22–24].
2. Evaluation Criterion: The degree of correlation between muscle activation as indicated by the MMG signal and the corresponding sEMG signal served as the metric for assessing the efficacy of the MMG signal extraction algorithm. A greater correlation coefficient signifies enhanced performance. Furthermore, Envelope Entropy serves as a metric for quantifying the irregularity inherent in sub-signals derived through various signal extraction methodologies. This metric facilitates the comparative assessment of energy levels associated with irregular noise components within the sub-signal spectra. A diminished lower envelope entropy signifies a reduction in noise, thereby correlating with an enhancement in signal quality.

Lower Limb Motion Classification

1. SCNGO-TCN-Attention Model: SCNGO Optimization Algorithm: Methodology: The Sine-Cosine Northern Goshawk Optimization (SCNGO) algorithm represents a refined iteration of the Northern Goshawk Optimization (NGO) algorithm, integrating sine and cosine functions along with refraction backward learning mechanisms. The position updating mechanism of the conventional NGO algorithm has been refined to augment both its exploratory and exploitative performance [25]. In the present study, the SCNGO algorithm is employed to optimize three pivotal parameters of the Temporal Convolutional Network with Attention (TCN-Attention) model, namely the learning rate, convolution kernel size, and the number of neurons. The primary optimization objective is to enhance the accuracy of the test dataset to its maximum potential. TCN-Attention Model: Methodology: The TCN-Attention model integrates the robust features of Temporal Convolutional Networks (TCNs) with the efficacy of attention mechanisms. Temporal Convolutional Networks (TCNs) are renowned for their inherent parallelizability and computational efficiency, rendering them particularly adept at handling sequential data processing tasks. The attention mechanism facilitates the model’s capacity to selectively concentrate on pertinent segments of the input sequence, thereby augmenting its proficiency in discerning long-range dependencies and salient features. The hybrid model exhibits significant utility in scenarios necessitating the concurrent capture of temporal dependencies and the selective emphasis on specific features, thereby rendering it highly appropriate for the classification of lower limb motion [26–29].
2. Kernel Principal Component Analysis (KPCA): Methodology: Kernel Principal Component Analysis (KPCA) is employed to distill pivotal information from the feature set, concurrently eliminating superfluous data [30]. This procedure enhances the model’s training efficiency by diminishing the dimensionality of the feature space, thereby preserving the most pertinent information. Advantages: Kernel Principal Component Analysis (KPCA) optimizes the model’s efficacy by streamlining the feature set, thereby facilitating expedited training durations and potentially augmenting generalization capabilities.

## 3. Experiments and Methods

This section primarily delineates the specifics of the lower limb exercise experimental protocol, the methodology for extracting mechanomyography (MMG) signals, the techniques employed for feature processing of MMG data, and the approach utilized for classifying lower limb activities. The experiment starts on January 2, 2024 and ends on January 5, 2024. From January 6 to October 15, 2024, the data were accessed for research purposes and authors had access to information that could identify individual participants during or after data collection.

### 3.1. Experimental Procedures

To obtain the necessary data, a sequence of rigorously controlled experiments was executed, involving a cohort of thirty participants, all within the age range of 24 to 28 years. Subjects were directed to attach the Cometa wireless myoelectricity apparatus, a state-of-the-art wireless surface electromyography (sEMG) acquisition system produced by Cometa, Italy, to their right lower limb. The position of sensors was shown in figure 1. The apparatus is outfitted with four 9-axis inertial measurement units (IMUs) and four surface electromyography (sEMG) electrodes. The apparatus was employed to collect triaxial acceleration data, which is crucial for the acquisition of mechanomyography (MMG) and surface electromyography (sEMG) signals. This investigation encompassed six discrete lower limb activities: standing, sitting, recumbency, ambulation, stair descent, and stair ascent. The experiments were conducted within a controlled laboratory environment, wherein participants were instructed to perform the activity sequence in a manner that emulated their typical behavioral patterns. The detailed parameters and configuration of the experimental protocol are outlined in Table 1.

**Figure. 1.**
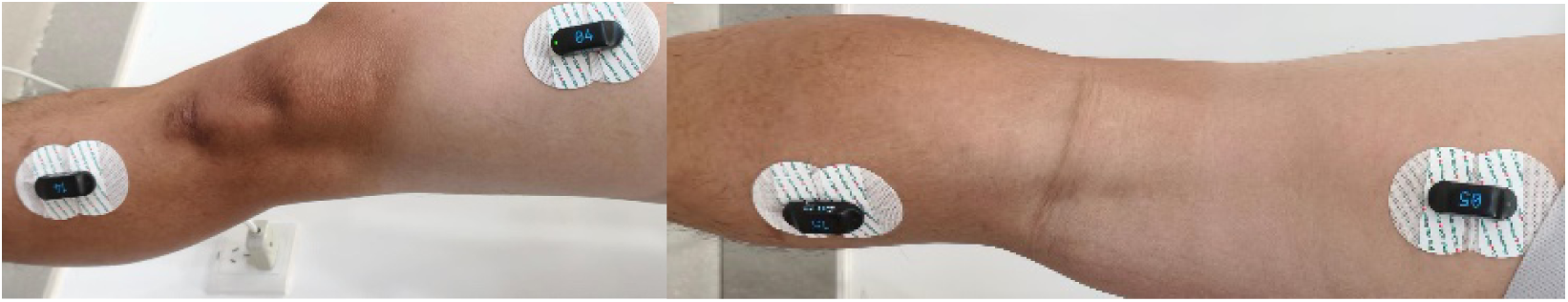
Position of sensors.

**Table 1.**
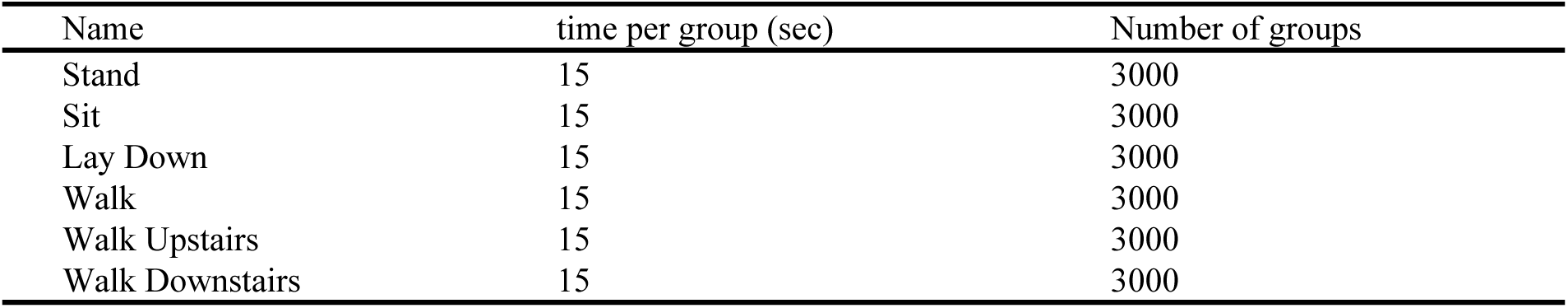
List of experiment.

### 3.2. Data Acquisition

The acceleration and surface electromyography (sEMG) signals were recorded at a sampling frequency of 2000 Hz. Subsequently, the signals were partitioned into discrete, fixed-width sliding windows, each encompassing a duration of 3 seconds, with an inter-window overlap of 50%. To account for the integration of mechanomyography (MMG) signals originating from the subjacent musculature within the three-axis acceleration data, the root mean square (RMS) of the triaxial acceleration was calculated as a preliminary step preceding the extraction of the MMG signals. Subsequently, Feature Mode Decomposition (FMD) was employed to deconstruct the four-channel acceleration signal, which encompassed the three-axis acceleration data and the Root Mean Square (RMS), thereby isolating the respective Mechanomyography (MMG) signals. Utilizing this methodology, a comprehensive dataset comprising 16 channel mechanomyogram (MMG) signals was generated, with the pertinent details delineated in Table 2

**Table 2.** Channel of MMG signals.

| Name | Number of channel |
| --- | --- |
| MMG of hamstrings | 4 |
| MMG of rectus femoris | 4 |
| MMG of tibialis anterior | 4 |
| MMG of gastrocnemius | 4 |

### 3.3. Data Processing

#### 3.3.1 Extraction of MMG based on Feature Mode Decomposition

In the present study, Feature Mode Decomposition (FMD) was utilized for the extraction of Mechanomyogram (MMG) signals. The proposed Finite Mode Decomposition (FMD) technique is fundamentally designed to decompose complex signals into discrete modes by employing adaptively configured finite-impulse response (FIR) filters. Utilizing the benefits of correlated kurtosis, FMD adeptly integrates the assessment of both impulsiveness and periodicity inherent in MMG signals simultaneously. A Finite Impulse Response (FIR) filter bank, initialized with a Hanning window, is first established to facilitate the decomposition process. Following this, an estimation and updating mechanism for the period is deployed to precisely detect and synchronize with the MMG signals. In the concluding phase, the mode selection process entails the exclusion of redundant and admixed modes to guarantee the integrity and precision of the isolated MMG signals. The extraction process was shown in figure 2.

**Figure. 2.**
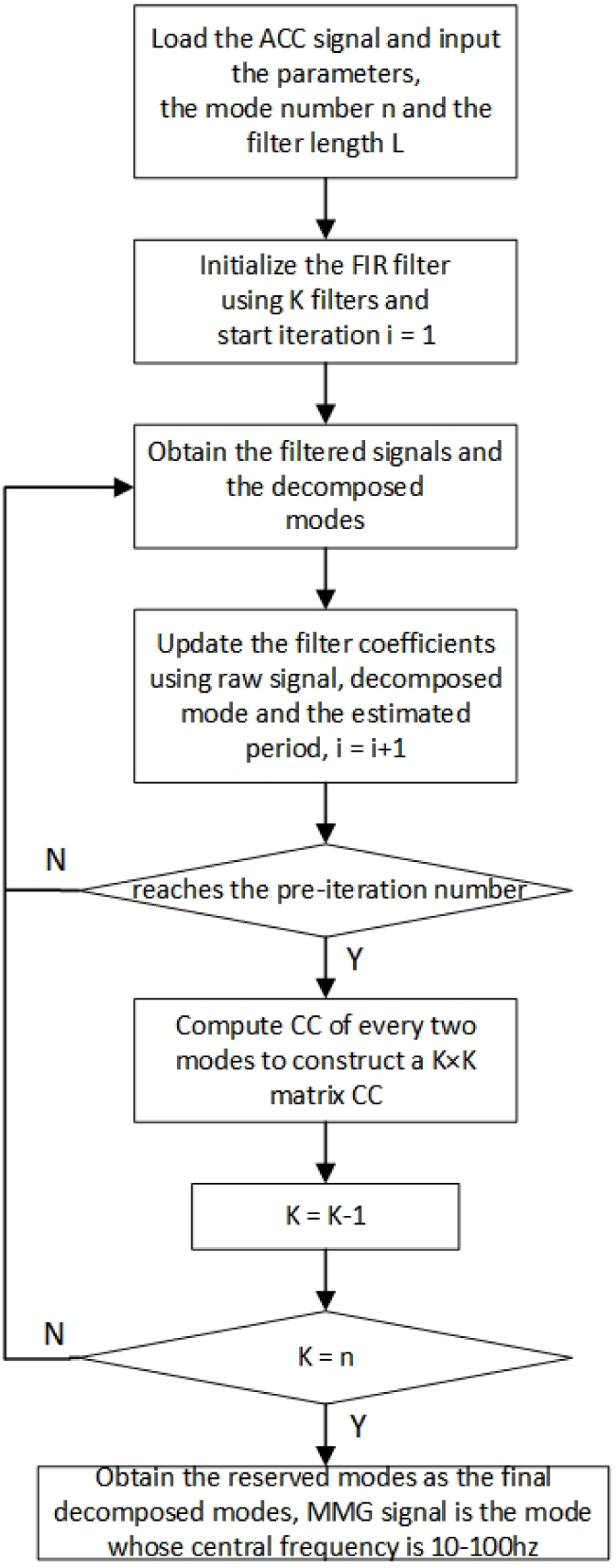
MMG extraction process based on FMD.

#### 3.3.2 Calculating of MMG features

In the present study, feature vectors corresponding to each MMG signal were extracted from the aforementioned sampling windows. Table 3 delineates the comprehensive array of metrics employed for the analysis of signals across both temporal and spectral domains. A robust dataset comprising 448 distinct features was meticulously compiled to comprehensively characterize each active window. To enhance the assessment of model efficacy, the dataset was systematically divided into two distinct subsets through random partitioning: a proportion of 70% was designated for training purposes, whereas the residual 30% was allocated for testing.

**Table 3.** List of features.

| Feature Function | Description |
| --- | --- |
| mean | Mean value |
| std | Standard deviation |
| mad | Median absolute value |
| max | Largest values in array |
| min | Smallest value in array |
| energy | Average sum of the squares |
| iqr | Interquartile range |
| entropy | Signal Entropy |
| arCoeff | Autocorrelation coefficients |
| correlation | Correlation coefficient |
| maxFreqInd | Largest frequency component |
| meanFreq | Frequency signal weighted average |
| skewness | Frequency signal Skewness |
| kurtosis | Frequency signal Kurtosis |

#### 3.3.3 dimensionality reduction of MMG features

In the present study, Kernel Principal Component Analysis (KPCA) was utilized for the purpose of dimensionality reduction of Mechanomyogram (MMG) feature sets. Kernel Principal Component Analysis (KPCA) represents an advanced machine learning methodology specifically tailored for nonlinear dimensionality reduction. This technique extends the functional scope of the traditional Principal Component Analysis (PCA), a linear approach that primarily focuses on the identification and extraction of the most prominent features or principal components within a given dataset. By projecting the initial dataset into an elevated-dimensional feature domain via a nonlinear transformation governed by a kernel function, Kernel Principal Component Analysis (KPCA) facilitates the calculation of dot products within this domain, obviating the necessity for explicit computation of the transformed dataset. The implicit computation enables the projection of data onto the principal components within the feature space, obviating the need for direct computation of these components. Kernel Principal Component Analysis (KPCA) exhibits significant utility in addressing datasets characterized by non-linear separability, thereby augmenting the efficacy of ensuing analytical processes or machine learning algorithms. The process of KPCA was shown in figure 3.

**Figure. 3.**
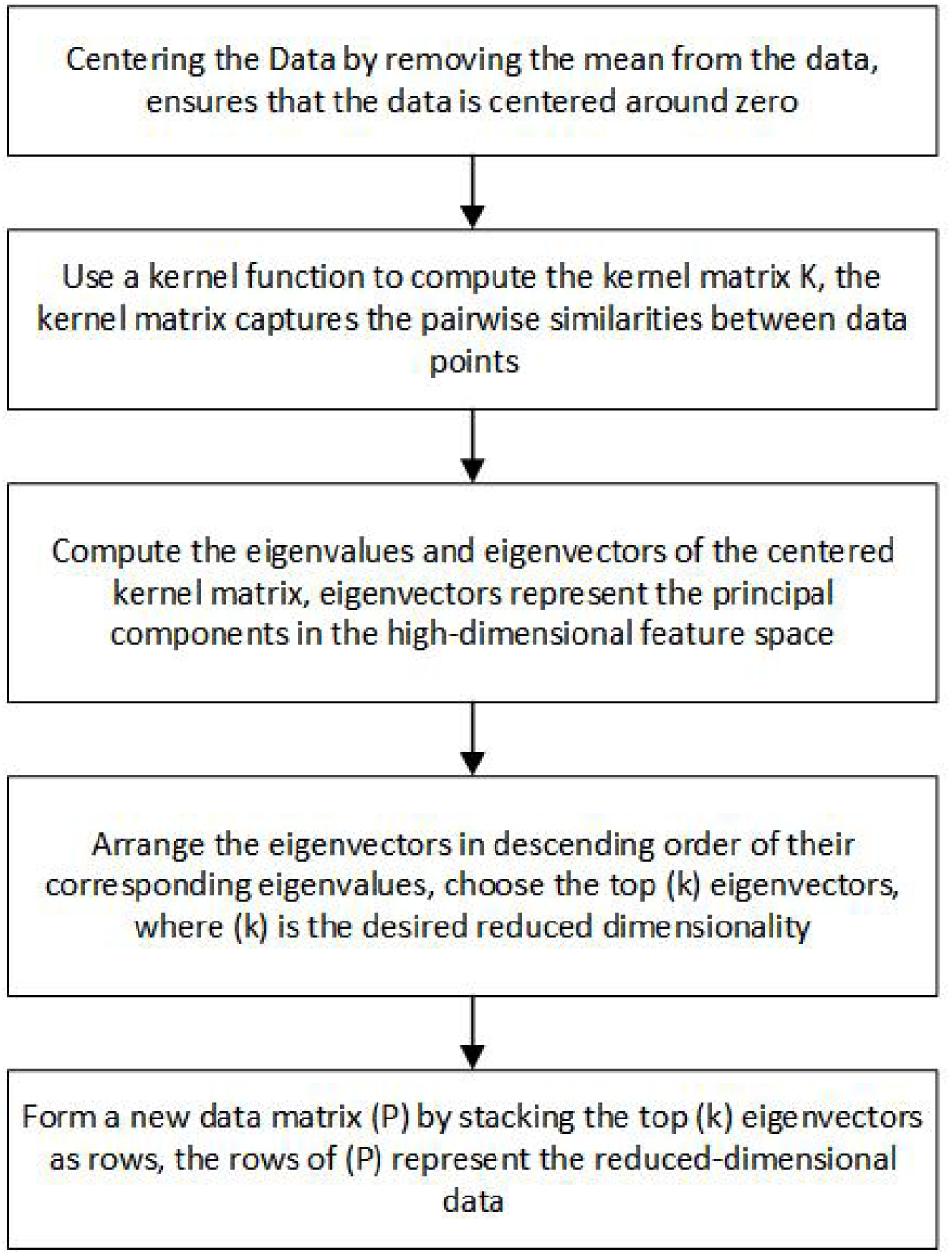
processing of KPCA.

### 3.4. Lower-Limb Activity Classification Methods

In the present study, a novel SCNGO-TCN-Attention classification algorithm was introduced to facilitate the accurate categorization of lower-limb activities. The algorithm was enhanced through the integration of Temporal Convolutional Networks (TCNs), an attention mechanism, and the refined Northern Goshawk Optimization Algorithm (SCNGO). Subsequent sections will delineate the attributes and interrelationships of the constituent elements. The overall processes was shown in figure 4.

**Figure. 4.**
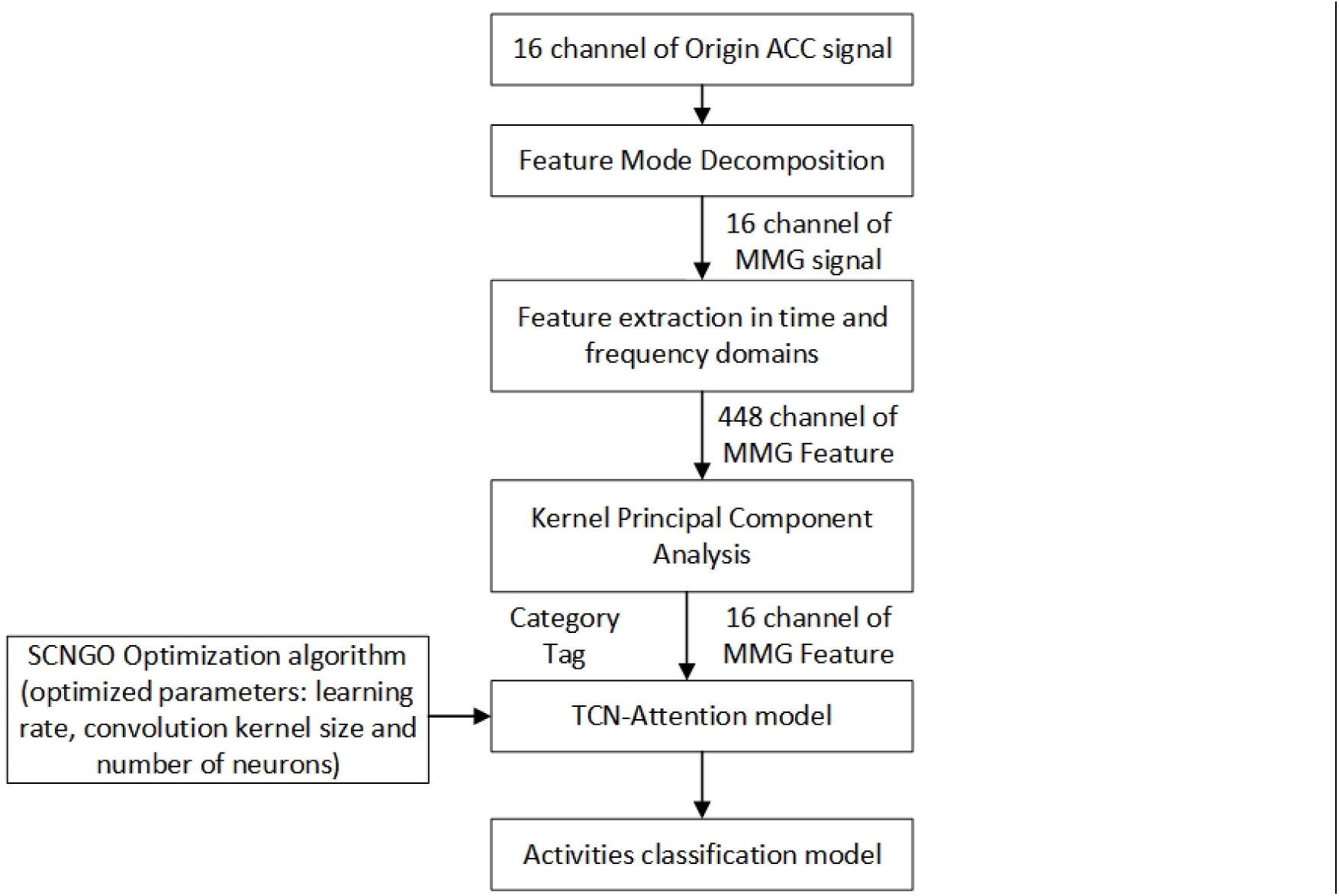
overall processes of Lower Limb Activities Classification Method.

#### 3.4.1 TCN-Attention model

Temporal Convolutional Networks (TCNs), when integrated with Multi-Head Attention mechanisms, constitute a robust methodology for the analysis of sequential data, effectively amalgamating the inherent advantages of convolutional neural networks (CNNs) and attention-based frameworks. The following text serves as an introductory exposition of the model under consideration:

Temporal Convolutional Networks (TCNs)

Temporal Convolutional Networks (TCNs) represent a specialized adaptation of Convolutional Neural Networks (CNNs) that are meticulously engineered to effectively process temporal data. In contrast to conventional recurrent neural networks (RNNs) and long short-term memory (LSTM) networks, which sequentially process sequences element by element, temporal convolutional networks (TCNs) concurrently analyze the entire sequence. The parallel processing capacity inherent in Temporal Convolutional Networks (TCNs) significantly enhances their training efficiency and accelerates the learning process, particularly when dealing with extensive sequence data.

Key Features of TCNs:

1. Temporal Convolutional Networks (TCNs) leverage dilated convolutions to expand the receptive field while maintaining a minimal increase in the parameter count. Dilated convolutions incorporate intervals between filter elements, thereby enabling the network to apprehend dependencies across extended temporal intervals.
2. Causal Convolutions: To guarantee that the output at any given time step is influenced exclusively by past and current inputs, thereby excluding any dependency on future inputs, Temporal Convolutional Networks (TCNs) utilize causal convolutions. This attribute is pivotal for real-time applications, guaranteeing that the model adheres to the temporal sequence of the data.
3. Residual Connections: Drawing inspiration from the ResNet architecture, residual connections serve to alleviate the vanishing gradient issue, thereby enabling the network to effectively learn and utilize deeper architectural configurations. These interconnections facilitate the propagation of gradients across the network, thereby enhancing the efficiency of training processes and promoting more effective convergence.

Multi-Head Attention Mechanism

The Multi-Head Attention mechanism constitutes a pivotal element within the Transformer architecture, initially developed for applications in natural language processing. The architecture facilitates the model’s capacity to concurrently focus on disparate segments of the input sequence, thereby encapsulating intricate dependencies and interrelationships.

Key Features of Multi-Head Attention:

1. Self-Attention Mechanism: The self-attention mechanism facilitates the capacity of each position within a sequence to engage with all positions in the preceding layer. This interaction permits the model to dynamically assign varying degrees of significance to distinct segments of the input data.
2. Utilization of Multiple Attention Heads: The deployment of multiple attention heads enables the model to concurrently concentrate on diverse facets of the input sequence, thereby enhancing its capacity for comprehensive analysis. Each head independently acquires the capacity to focus on distinct patterns, thereby enriching the overall representation of the input data.
3. Linear Transformations: Input embeddings undergo linear projection to generate query, key, and value vectors. The attention mechanism calculates the weighted summation of value vectors, determined by the degree of similarity between the query and key vectors.

Combining TCNs and Multi-Head Attention

The integration of Temporal Convolutional Networks (TCNs) and Multi-Head Attention mechanisms synergistically harnesses the respective advantages of these methodologies, thereby fostering the development of a robust framework for the efficient processing of sequential data.

1. Temporal Dynamics: Temporal Convolutional Networks (TCNs) effectively encapsulate the temporal dynamics of sequences via dilated and causal convolutional mechanisms, thereby facilitating the efficient management of long-range dependencies within the data.
2. Attention Mechanisms: The implementation of Multi-Head Attention facilitates the model’s ability to selectively concentrate on pertinent segments of the sequence, thereby effectively capturing intricate interactions and dependencies that may otherwise remain unexplored by solely convolutional methodologies.
3. Parallel Processing: The inherent parallelism of Temporal Convolutional Networks (TCNs) and attention mechanisms significantly enhances the model’s scalability and efficiency, rendering it particularly well-suited for real-time and large-scale application scenarios.

#### 3.4.3 The improved Northern Goshawk Optimization Algorithm

The Enhanced Northern Goshawk Optimization Algorithm (SCNGO) was developed as an extension of the Northern Goshawk Optimization Algorithm (NGO), originally proposed by MOHAMMADDEHGHANI and formally released in 2022. This algorithm emulates the hunting behavior of the northern goshawk, encompassing stages such as prey identification and assault, pursuit, and evasion. The refinement strategy pertains to the Sparrow Optimization Algorithm, with the following enhancements delineated:

1. The refraction-based reverse learning strategy is employed to initialize the individuals within the Northern Goshawk optimization algorithm. The fundamental concept entails broadening the search horizon by computing the inverse of the existing solution, thereby facilitating the identification of a superior alternative for the specified problem.
2. The conventional position update mechanism of the Goshawk algorithm has been supplanted by a sine-cosine optimization strategy.
3. The step search factor within the sine-cosine strategy has been refined. The initial step size search factor exhibited a linear decrement, which was suboptimal for achieving a balanced enhancement of both global search and local refinement capabilities in the Northern Goshawk optimization algorithm.

## 4. Results

### 4.1. Results of MMG extraction

In the present study, a novel methodology termed Feature Mode Decomposition (FMD) was introduced for the precise extraction of Mechanomyogram (MMG) signals from accelerometer-derived acceleration data. Figures 5 and 6 respectively depict the extraction outcomes within the temporal and spectral domains. Considering that MMG signals generally reside within the 10-100 Hz frequency spectrum, the data presented in these figures unequivocally demonstrate that FMD effectively isolates MMG signals exhibiting frequencies within this specified range. Nonetheless, spectral analysis alone does not comprehensively capture the capability of Frequency Modulation Detection (FMD) to mitigate random noise.

**Figure. 5.**
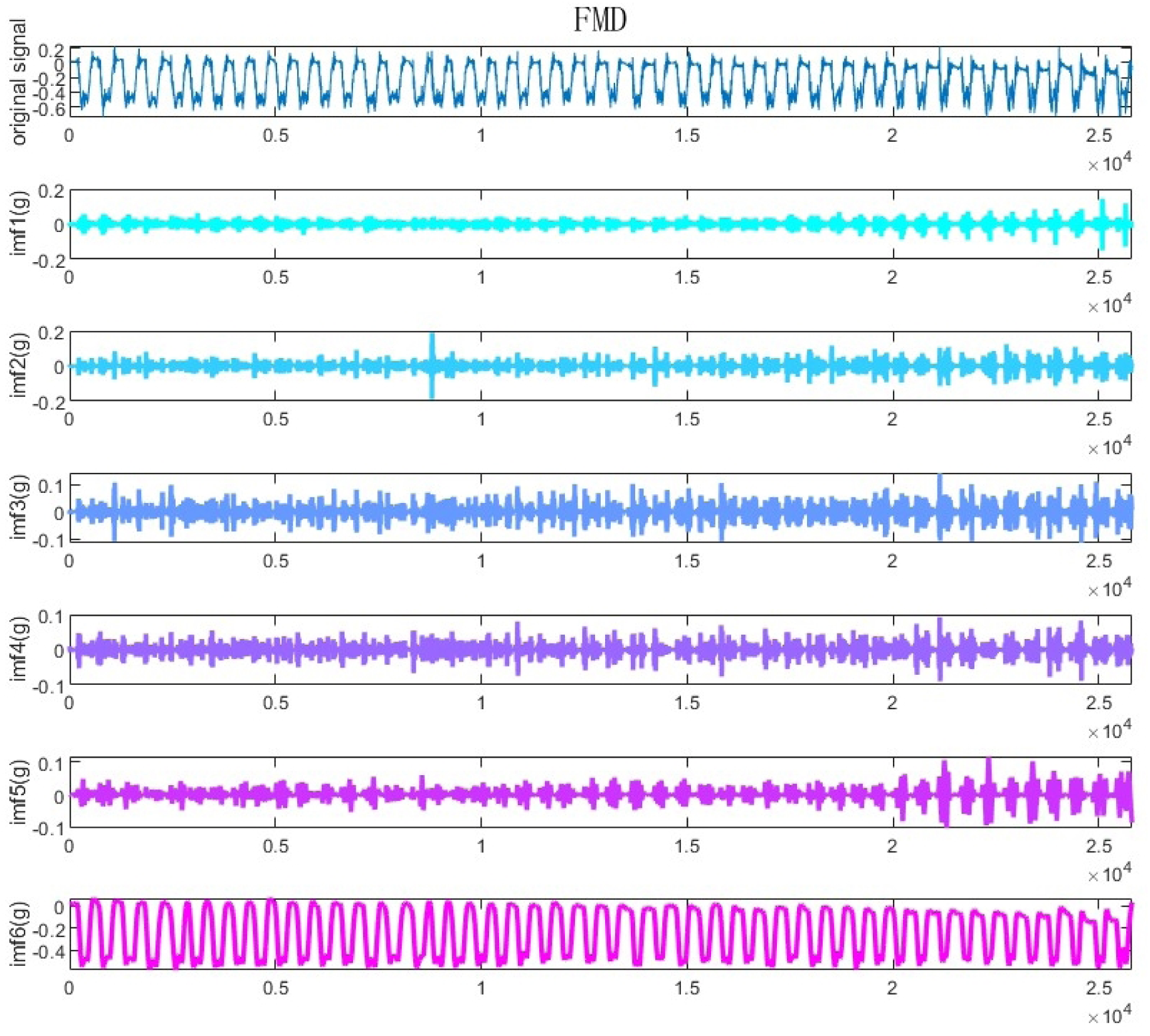
extraction result of FMD in time domain.

**Figure. 6.**
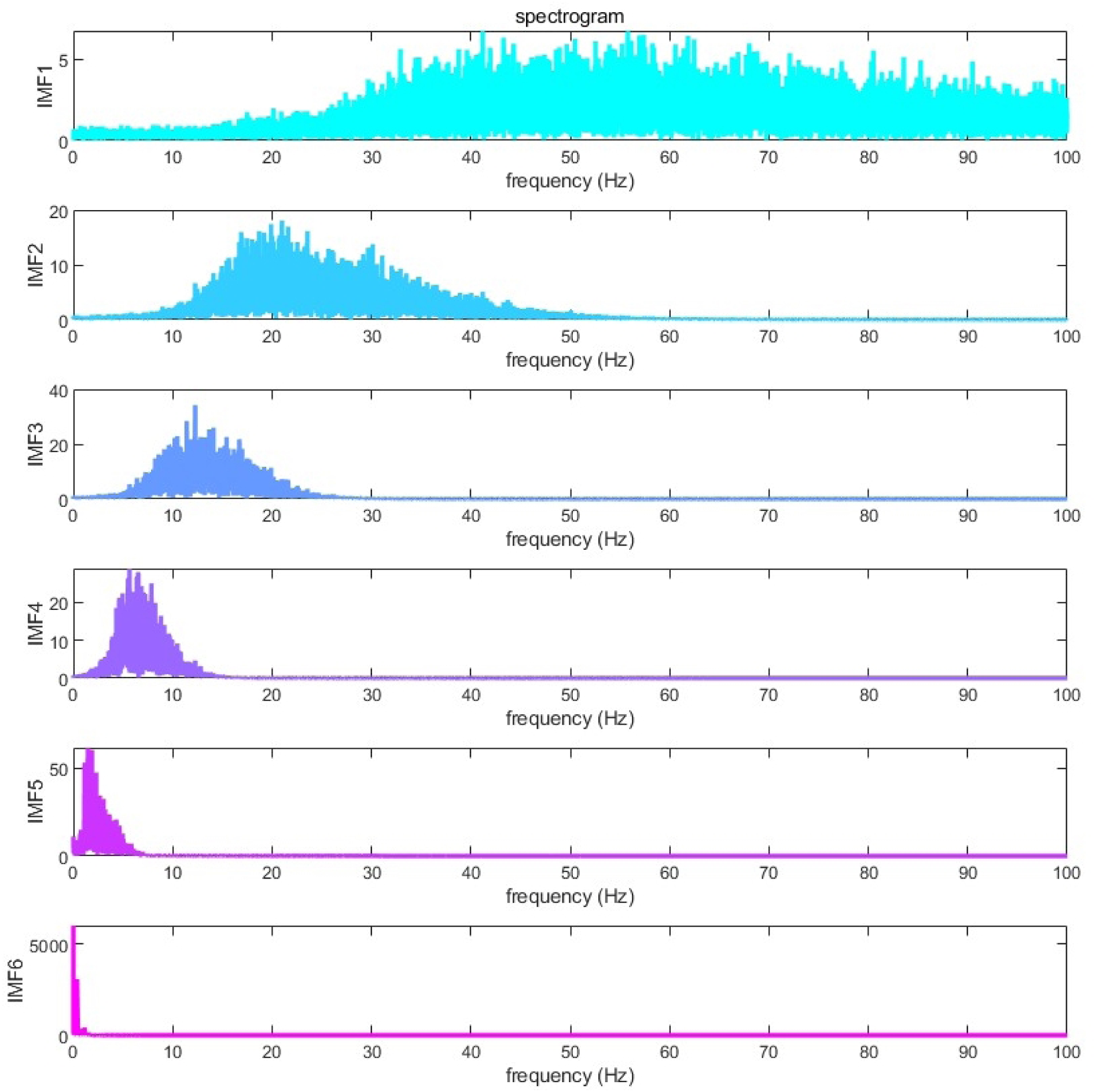
extraction result of FMD in frequency domain.

To assess the efficacy of noise suppression, the envelope entropy of the FMD and various other signal decomposition algorithms was quantified. As depicted in Figure 7, the envelope entropy of the FMD exhibited the lowest value, recorded at 8.13.This evidence demonstrates that FMD possesses enhanced efficacy in mitigating random noise, thereby augmenting the quality and dependability of the extracted MMG signals. The reduced lower envelope entropy indicates that FMD offers a more pristine and resilient representation of MMG signals relative to alternative decomposition techniques..

**Figure. 7.**
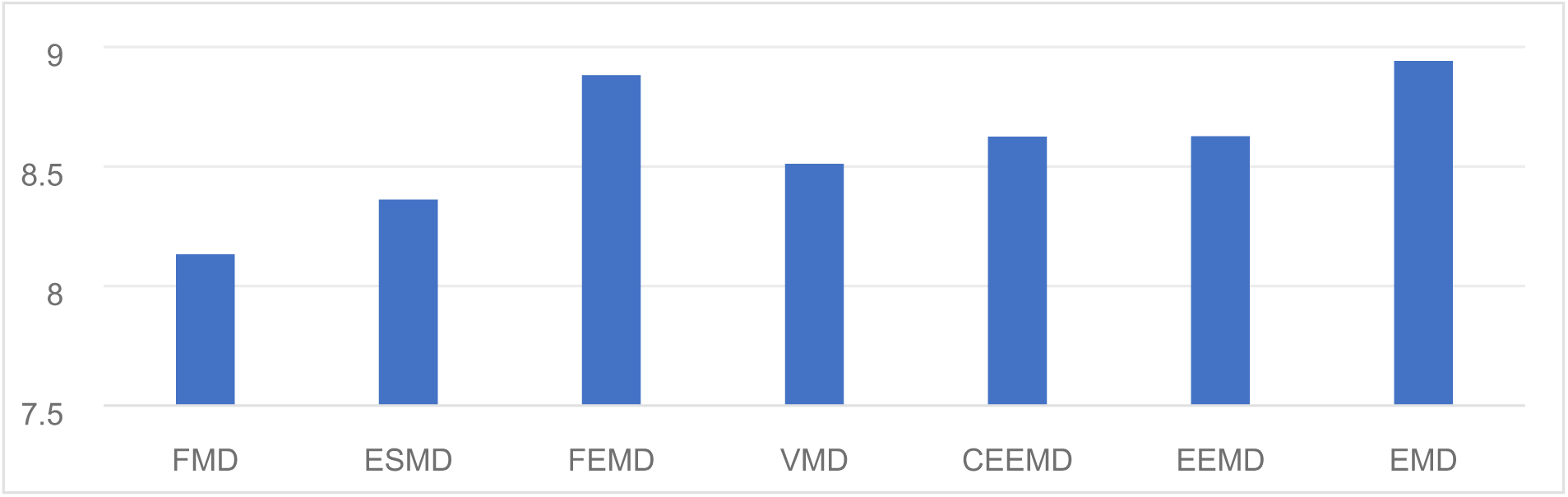
Envelope entropy of different algorithm.

In this study, we conducted an analysis of both mechanomyography (MMG) and surface electromyography (sEMG) signals, which are produced as a result of muscle contraction. MMG specifically denotes the mechanical vibration data, whereas sEMG records the electrical activity. Considering their common derivation, it is plausible to authenticate the extracted MMG signal by assessing the correlation between MMG and sEMG. Given that both MMG and sEMG represent non-stationary time series data, a sequence of pre-processing procedures was imperative prior to the execution of correlation analysis. The pre-processing procedures encompassed the following stages:

1. Extraction of MMG from Raw Acceleration Data: The raw acceleration dataset was systematically processed to isolate the MMG signal through the application of the novel Feature Mode Decomposition (FMD) technique.
2. DC Offset Correction: MMG and sEMG signals were systematically adjusted to eliminate any existing DC offset, thereby ensuring that the signals were accurately centered at the zero baseline.
3. Application of Butterworth Bandpass Filtering: To isolate the pertinent frequency ranges, a Butterworth bandpass filter was employed for both mechanomyographic (MMG) and surface electromyographic (sEMG) signals, with the frequency bands set to 10-100 Hz for MMG and 20-450 Hz for sEMG, respectively.
4. Full-Wave Rectification Process: The implementation of full-wave rectification was applied to both MMG and sEMG signals, effectively transforming all negative values to positive, thereby accentuating the signal amplitudes.
5. Extraction of Linear Envelopes: The linear envelopes of both mechanomyographic (MMG) and surface electromyographic (sEMG) signals were derived to refine the rectified signals, thereby accentuating the overarching trends.
6. Normalization: To ensure a consistent scale and enable meaningful comparative analysis, both MMG and sEMG signals underwent normalization procedures.

Figure 8 illustrates that the correlation coefficient between the MMG and sEMG signals, derived via FMD extraction, reached the maximum value of 0.87.The data suggest that the MMG signals derived via the FMD technique exhibit the highest degree of correlation with the sEMG signals, thereby affirming the efficacy and precision of the FMD method in the extraction of MMG signals. The strong correlation observed indicates that the MMG signals extracted via the FMD method precisely correspond to the underlying muscle activity, as evidenced by the sEMG signals..

**Figure. 8.**
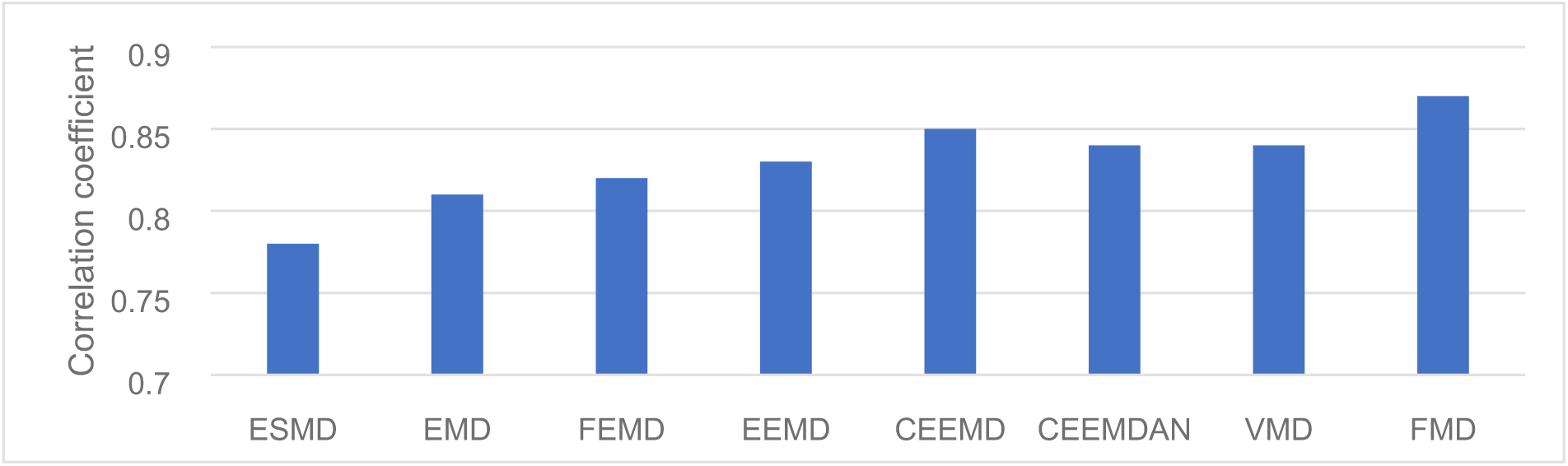
Correlation coefficient of different algorithm.

### 4.2. Results of feature dimensionality reduction

In the current investigation, Kernel Principal Component Analysis (KPCA) was introduced as a method for dimensionality reduction within the feature space, with Principal Component Analysis (PCA) employed as a comparative benchmark to assess the effectiveness of KPCA. The classification accuracy and computational time were determined by averaging the results across 100 iterations of the training and testing processes. As illustrated in Figure 9, an augmentation in the quantity of components resulted in a proportional elevation in the cumulative contribution, wherein the cumulative contribution of Kernel Principal Component Analysis (KPCA) consistently surpassed that of Principal Component Analysis (PCA).This evidence suggests that Kernel Principal Component Analysis (KPCA) exhibits superior efficacy in extracting salient information from principal components relative to Principal Component Analysis (PCA).

**Figure. 9.**
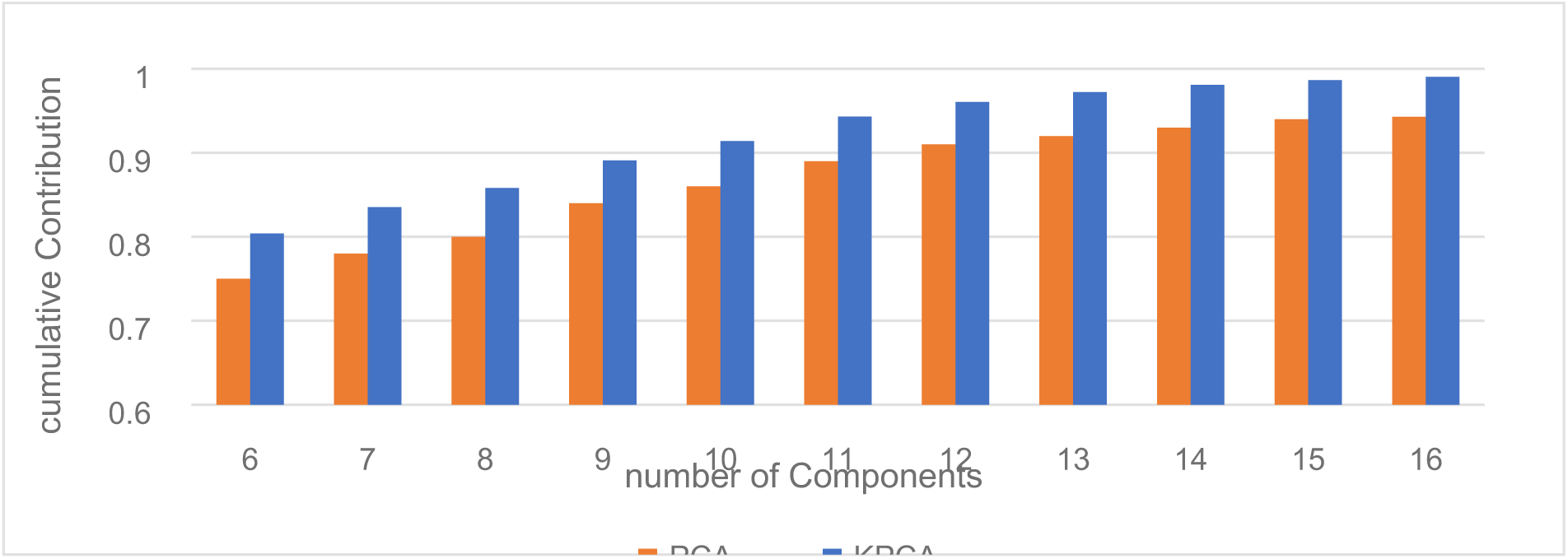
Cumulative Contribution of different feature dimensionality reduction methods.

Despite exhibiting enhanced efficacy in the extraction of salient features, Kernel Principal Component Analysis (KPCA) incurred a substantial computational overhead, being approximately quadruple that of Principal Component Analysis (PCA). Nevertheless, despite the constrained feature set, the mean computational duration for Kernel Principal Component Analysis (KPCA) remained comparatively low, averaging 1.21 seconds. This indicates that although Kernel Principal Component Analysis (KPCA) is computationally more demanding, its superior capability to capture non-linear relationships and extract significant features warrants the increased computational cost, particularly in contexts where the feature set is not excessively large.

The results of classification corresponding to different quantities of principal components are delineated in Table 4.The components were derived utilizing Kernel Principal Component Analysis (KPCA), followed by the application of the TCN-Attention mechanism for the ensuing classification process. The initial column of Table 4 delineates the quantity of components employed. It is demonstrably clear that Kernel Principal Component Analysis (KPCA) adeptly isolates prominent feature combinations from high-dimensional datasets, thereby substantially augmenting the training efficiency of the classification model while maintaining classification accuracy with negligible degradation.

**Table 4.** result of different number of components.

| Number of components | Training time(s) | Classification time(ms) | Accuracy |
| --- | --- | --- | --- |
| 6 | 13.1 | 11.3 | 92.6% |
| 7 | 13.5 | 12.1 | 93.1% |
| 8 | 13.9 | 11.5 | 93.5% |
| 9 | 14.9 | 12.5 | 94.0% |
| 10 | 15.8 | 12.4 | 94.2% |
| 11 | 17.4 | 12.6 | 94.4% |
| 12 | 18.6 | 13.1 | 95.6% |
| 13 | 19.5 | 12.9 | 96.3% |
| 14 | 20.9 | 13.5 | 97.0% |
| 15 | 22.8 | 14.2 | 97.1% |
| 16 | 24.9 | 14.6 | 97.2% |
| 448 | 589.7 | 98.6 | 97.8% |

### 4.3. Results of hyperparameters optimization based on SCNGO algorithm

To assess the influence of hyperparameter optimization via the SCNGO algorithm, a comparative evaluation was performed between the baseline classification model and the refined, optimized model.Prior to the initiation of training, a total of 16 principal components were derived through the application of Kernel Principal Component Analysis (KPCA).The confusion matrix and the fitness curve are illustrated in Figures 10 and 11, respectively.The data presented in these figures demonstrate that by the fourth generation, the loss function achieved convergence, with the error rate being reduced to 0.016.The outcomes of the parameter optimization process are succinctly encapsulated in Table 5. Additionally, a comparative analysis of various optimization algorithms, encompassing SCNGO, is delineated in Table 6.An examination of the data presented in Tables 5 and 6 indicates that SCNGO exhibits superior performance compared to NGO, with notable enhancements observed in both accuracy and training efficiency.

**Figure. 10.**
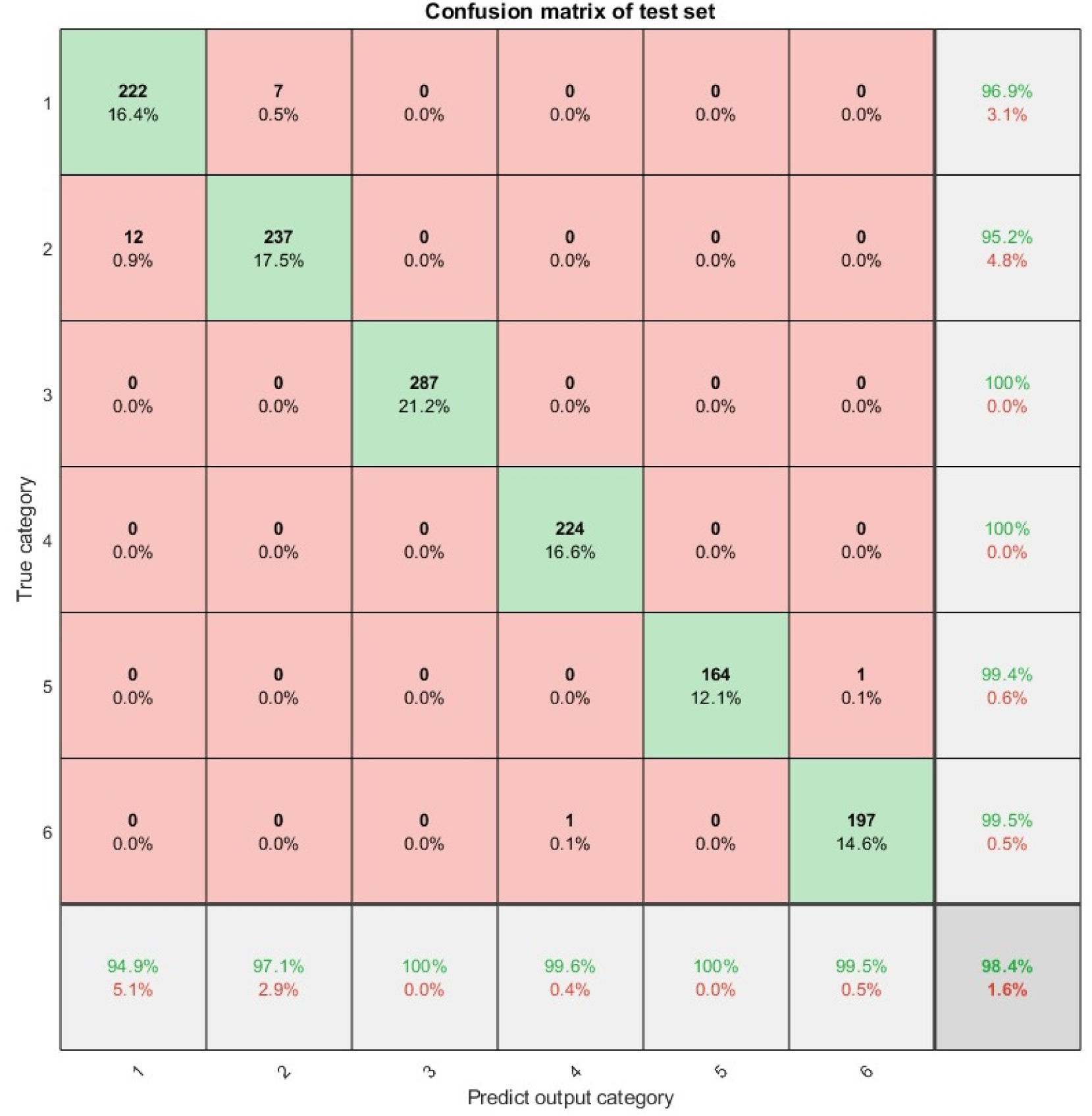
classification result of SCNGO-TCN-Attention model.

**Figure. 12.**
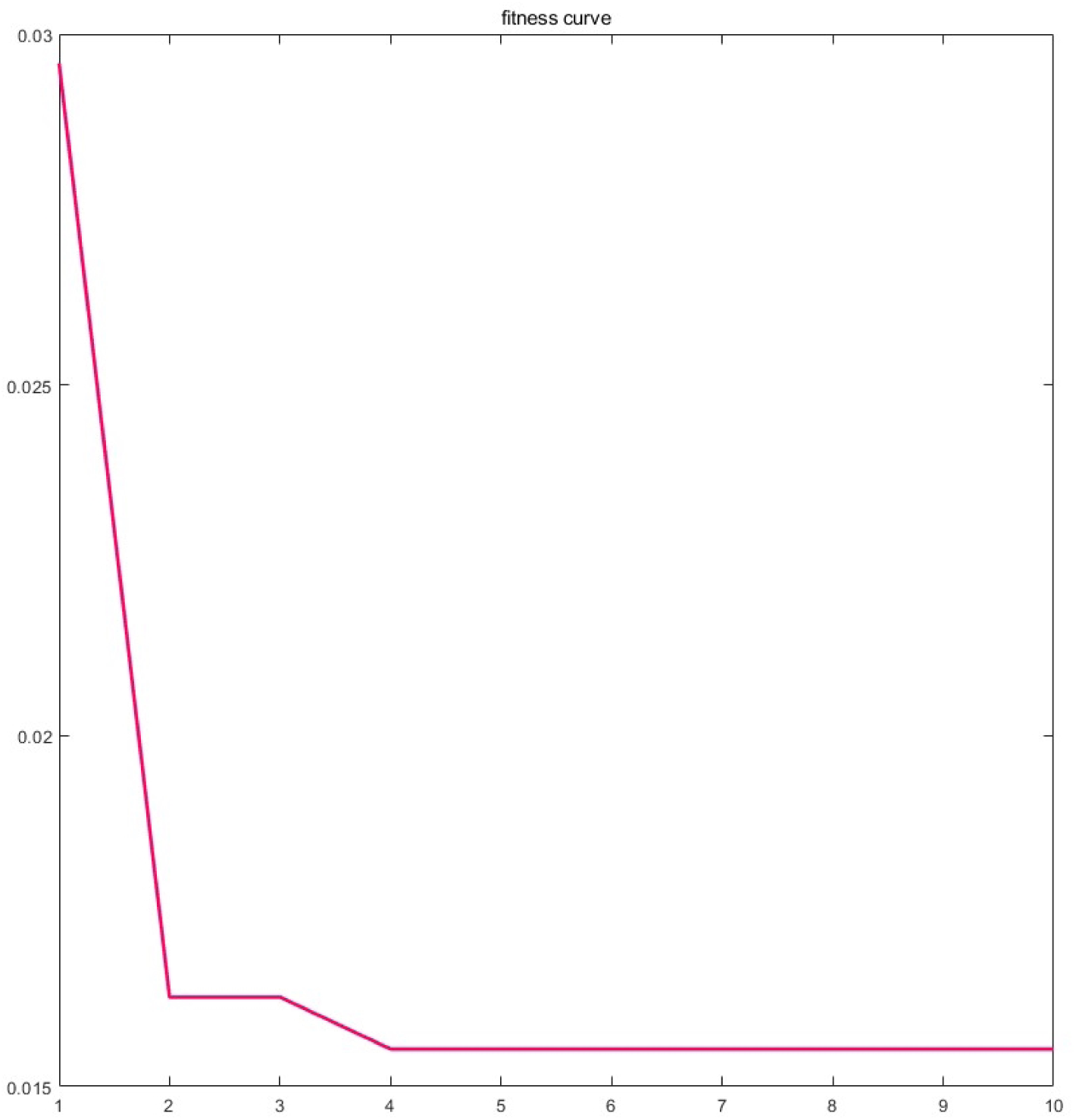
result of fitness curve.

**Table 5.** parameters optimization result.

| parameters | value |
| --- | --- |
| Learning rate | 0.0097 |
| Number of convolution kernels | 3 |
| Convolution kernel size | 9 |
| Number of residual blocks | 3 |
| self-attention layers | 4 |
| Number of self-attention heads | 256 |

**Table 6.** comparison of NGO-TCN-Attention and SCNGO-TCN-Attention.

| algorithm | Training time(s) | Classification time(ms) | Accuracy |
| --- | --- | --- | --- |
| TCN-Attention | 24.9 | 14.6 | 97.2 |
| NGO-TCN-Attention | 1032.7 | 14.5 | 97.4 |
| SCNGO-TCN-Attention | 954.5 | 14.4 | 98.4 |

### 4.4. Comparison of different classification algorithm

To evaluate the efficacy of the proposed SCNGO-TCN-Attention algorithm with respect to training accuracy, computational speed, and classification precision, a comparative analytical study was performed utilizing multiple benchmark classification algorithms. Prior to initiating model training, a total of 16 principal components were derived through the application of Kernel Principal Component Analysis (KPCA). The comparative outcomes are succinctly presented in Table 7. Based on the data presented in Table 7, the SCNGO-TCN-Attention algorithm demonstrated the highest level of accuracy compared to the other methodologies examined. The training and classification durations associated with the proposed method were observed to occupy an intermediate position, suggesting that these temporal metrics fall within acceptable thresholds.

**Table 7.** comparison of different algorithms.

| algorithm | Training time(s) | Classification time(ms) | Accuracy |
| --- | --- | --- | --- |
| TCN-Attention | 24.9 | 14.6 | 97.2 |
| SCNGO-TCN-Attention | 954.5 | 14.4 | 98.4 |
| BP | 126 | 5.5 | 91.5 |
| LSTM | 239 | 11.5 | 96.2 |
| BiLSTM | 395 | 12.1 | 96.8 |
| CNN | 951 | 5.3 | 94.6 |
| TCN | 894 | 6.1 | 95.1 |
| KNN | 1068 | 28.9 | 95.6 |
| SVM | 2657 | 31.6 | 94.6 |
| RF | 95 | 19.4 | 93.5 |
| RBF | 784 | 39.1 | 94.8 |
| ELM | 815 | 25.8 | 90.6 |

## 5. Conclusion and Discussion

The objective of this investigation was to mitigate the difficulties associated with the extraction of Mechanomyography (MMG) signals from unprocessed acceleration data and to develop a method for the accurate classification of human lower limb activities utilizing these extracted signals. The Feature Mode Decomposition (FMD) algorithm was utilized to isolate the MMG signals. Subsequently, surface Electromyography (sEMG) signals were employed for correlation analysis to ascertain the efficacy of the extracted MMG signals. The findings revealed that FMD not only efficiently mitigated random noise, as evidenced by the minimal envelope entropy value of 8.13, but also retained the utmost muscle contraction information, as indicated by a substantial correlation coefficient of 0.87 between the MMG and sEMG signals. The data indicate that FMD exhibits a high degree of efficacy in the extraction of MMG signals.

To augment the classification performance, a comprehensive feature extraction protocol was implemented, resulting in the derivation of 448 unique features from multi-channel MMG signals. Subsequently, Kernel Principal Component Analysis (KPCA) was employed to diminish the dimensionality of the extracted features. This process was succeeded by the initialization and training of a hybrid model SCNGO-TCN-Attention.The SCNGO-TCN-Attention model demonstrated an exceptional classification accuracy of 98.44%, surpassing the performance of conventional machine learning algorithms, including Backpropagation (BP), Long Short-Term Memory (LSTM), Bidirectional LSTM (BiLSTM), Convolutional Neural Networks (CNN), and additional methodologies. The findings underscore the robustness and efficacy of the proposed hybrid model in addressing the intricacies associated with complex time-series data.

## 6. Discussion

1. Advanced Methodology: The introduced technique presents a pioneering solution for precise human activity recognition, specifically designed for contexts where the utilization of costly apparatus and the availability of regulated settings are impractical. This renders the technology apt for practical applications in domains such as health surveillance, rehabilitative processes, and individual fitness monitoring.
2. Limitations and Prospects for Future Research: Although the introduced model exhibits superior performance metrics, its generalizability and applicability across diverse individual populations necessitate additional empirical validation. Subsequent studies should endeavor to augment both the sample size and its demographic heterogeneity, thereby fortifying the model’s robustness and broadening its applicability across diverse populations. Furthermore, the investigation of sophisticated signal processing methodologies and advanced deep learning frameworks holds the potential to significantly enhance the fidelity of MMG signals, thereby augmenting their utility in the domain of human activity recognition.
3. Computational Efficiency: While Kernel Principal Component Analysis (KPCA) demonstrated enhanced capability in extracting salient features, it incurred a greater computational expenditure relative to Principal Component Analysis (PCA). Nevertheless, the computational duration necessitated by Kernel Principal Component Analysis (KPCA) remained comparatively moderate, indicating that the equilibrium between computational expediency and the caliber of feature extraction is warranted, particularly when the feature set is not excessively voluminous. Future research endeavors may concentrate on enhancing the computational efficacy of Kernel Principal Component Analysis (KPCA) or investigating alternative methodologies for dimensionality reduction that achieve an optimal equilibrium between performance metrics and computational expenditure.
4. Real-World Applications: The SCNGO-TCN-Attention model demonstrates exceptional accuracy and robustness, rendering it a highly promising instrument for a diverse array of real-world applications. In the healthcare sector, this technology can be harnessed for patient monitoring and rehabilitation, thereby ensuring compliance with therapeutic protocols and bolstering recovery initiatives. In the realms of sports and fitness, this technology affords comprehensive insights into kinematic patterns, thereby facilitating the optimization of athletic performance.

In summary, this investigation presents a robust and efficacious methodology for the extraction and classification of human lower limb activities through the utilization of mechanomyography (MMG) signals. The integration of FMD for signal extraction, KPCA for feature dimensionality reduction, and the SCNGO-TCN-Attention model for classification demonstrates substantial promise in augmenting the precision and dependability of human activity recognition systems. This research establishes a robust groundwork for subsequent advancements within the domain, thereby facilitating the development of more advanced and dependable systems for human activity recognition.

## Acknowledgments

Thanks to the experiment participants and Postgraduate Research Practice Innovation Program of Jiangsu Province for their support of this study.

## Statements & Declarations

### Code or data availability

The code and data used during the current study are available from: https://gitee.com/baiyu1928/sc-tcn-at.

### Ethics approval

This work involved human subjects in its research. Approval of all ethical and experimental procedures and protocols was granted by the Medical Ethics Committee of Nanjing Medical University under approval number 2021-SR109.

### Consent to participate

Informed consent was obtained from all individual participants included in the study.

### Consent for publication

The authors affirm that human research participants provided informed consent for publication.

